## Supplementary Information for "A platform of patient-derived microtumors identifies therapeutic vulnerabilities in ovarian cancer"

SI Methods

**Viability measurement of OvCa PDM after isolation**

About n =10 PDM were removed from PDM culture into a 96-well clear bottom plate and were stained with 2 µM viability dye Calcein-AM (Invitrogen) and 5 µM SYTOX orange nucleic acid stain (Invitrogen) in 200 µl culture media. After 30 min incubation time, z-stack images of stained PDM were taken in FITC and TRITC channel with a spinning disk microscope (ZEISS CellObserver Z1). 2D images of 3D z-stack projections within the Zen 2.6 software were generated. Percentages of PDM viability were assessed by using the Imaris 8.0 software. 3D surface masks were created for FITC and TRITC channel. Thresholds were adjusted and the sum of the total volume of each surface mask was measured. To calculate the percentage of viable/dead cells, the volume of each channel was divided by the total volume and multiplied by 100.

**Histology/Immunohistochemistry**

For histology of ovarian cancer microtumors, n = 5-10 PDM were fixed for 1 hour in 4% Roti® Histofix (Carl Roth) at RT and then collected in a 40 µm cell strainer. PDM were incubated for 5 min in Harris Hematoxylin (Leica Biosystems), shortly washed in dH_2_O and dehydrated in an ethanol series (2x 50% ethanol, 2x 70% ethanol, each for 15 min). PDM were transferred into a Tissue-Tek® Cryomold® (Sakura) and embedded in Richard-Allan Scientific™ HistoGel™ (Thermo Fisher Scientific). After subsequent tissue processing with HistoCore PEARL (Leica Biosystems), PDM histogel-blocks were paraffin-embedded for sectioning. Three micrometer sections of FFPE PDM samples were prepared. Corresponding primary tumor tissue was received as cryosections (5-7 µm) from the Center for Women’s Health, Tuebingen. Primary tissue samples (OvCa #24-26) were directly taken from the delivered tumor sample, fixed in 4% Roti® Histofix overnight at RT and paraffin-embedded after tissue processing. For Hematoxylin and Eosin (H&E) staining, PDM-FFPE sections were baked (60 min, 55°C), deparaffinized and rehydrated in decreasing ethanol series. Afterwards the sections were stained with Harris Hematoxylin (Leica Biosystems) for 2-3 min. Sections were washed in tap water for 3 min, incubated in 2% acetic acid for up to 2 min, neutralized in 95% ethanol, stained with Eosin Y (alcoholic, Leica Biosystems) for 2 min and at last dehydrated by an increasing ethanol series ending with Roti® Histol (Carl Roth). H&E staining of corresponding primary tumor tissue samples (cryosections) was performed within the pathology department of the Women’s Health Clinic, Tuebingen.

For immunohistochemical staining of PDM-FFPE sections, antigen-retrieval was performed in TE buffer (10mM Tris/1mM EDTA, pH 9.0) at 95-98°C after slides were baked, deparaffinized and rehydrated in decreasing ethanol series to water. Cryosections of corresponding primary tumor tissue were simply rehydrated in PBS for 10 min. All sections (FFPE and cryosections) were subsequently incubated with BLOXALL® Endogenous Peroxidase and Alkaline Phosphatase Blocking Solution (VECTOR Laboratories) for 10 min. Slides were washed in PBS and incubated with blocking medium (PBST with 10% normal goat serum plus Novocastra™ Avidin) (Leica Biosystems) for 1 h at RT in a humidified chamber. After two washing steps in PBS, sections were incubated with primary antibodies (see SI Materials) diluted in PBS, 1% BSA and Novocastra™ Biotin (Leica Biosystems) overnight at 4°C in a humidified chamber. Slides were washed 2x in PBS and incubated with secondary antibodies (see SI Materials) in PBS plus 1% BSA (30 min, at RT). Slides were washed in PBS and incubated with R.T.U Horseradish Peroxidase Streptavidin (VECTOR laboratories) for 30 min. Staining was accomplished by using the DAB (Polymer) Kit Novocastra Novolink™ (Leica Biosystems) for 5 min. Slides were counterstained with hematoxylin QS (VECTOR Laboratories). After dehydration and clearance in a rising ethanol series to Roti® Histol, the slides were mounted with CV mounting medium (Leica Biosystems). For high quality staining of p53 and WT1 in PDM, immunohistochemistry stainings were performed by the Institute of Pathology and Neuropathology at the University Hospital Tuebingen using routine and standardized immunohistochemistry protocols. Stained FFPE/cryosections were imaged with Axio Scan Z1. All primary antibodies were validated in normal, healthy tissues as well as in FFPE and cryosections.

**Protein Profiling of PDM by RPPA**

To generate protein abundance profiles of PDM and to analyze their dynamic on/off-target changes upon treaments, Reverse Phase Protein Array (RPPA) protein profiling using Zeptosens technology (1, 2) was used. Cultured PDM (n = 100-200) were harvested, washed in HBSS, snap frozen in 0.65ml LoBind Eppis (Eppendorf) and stored at -80°C. To analyze treatment effects over time, four replicates of n = 25-35 PDM were treated with a drug, pooled, washed in HBSS and snap frozen after 30 min, 4 h and 72 h in 0.65 ml or 1.5 ml Protein LoBind tubes (Eppendorf) on dry ice. PDM pellets were stored at -80°C. For protein extraction from the low amount of PDM pellet, samples were lyophilized with a Epsilon 1-4 LSC plus instrument (Christ/Osterode). Lyophilisates were lysed with 6.25 µl CLB1 lysis buffer (Zeptosens) for 30 min (RT) at 1400 rpm (thermomixer) followed by centrifugation (5 min, 13200 rpm at RT). Protein amount was determined by Pierce™ Coomassie Plus™ (Bradford) Protein-Assay (Thermo Fisher Scientific). For printing, PDM lysates were adjusted to a uniform protein concentration (2 µg/µl) by diluting the samples in CLB1 if necessary, following vortexing, centrifugation (5 min, 13200 rpm, RT) and storage at -80°C. Adjusted PDM lysates were diluted 10-fold in RPPA spotting buffer CSBL1 and printed as replicate microarrays on Zeptosens hydrophobic chips (NMI TT) using a NanoPlotter 2 (GeSim), each sample at two technical replicates.. With the samples, fluorescence-labeled albumin (reference spots), internal standard lysates and further technical controls were co-printed for use as quality control. Freshly printed chips were blocked with 3% BSA solution, washed subsequently with dH_2_0, dried under a nitrogen stream and stored in the dark at 4°C.

Protein expression signals were measured using a direct two-step sequential fluorescence immunoassay. Up to six spotted chips (6x6 arrays) were assembled in a ZeptoCARRIER with fluidic structures. Well-characterized, pre-validated primary antibodies of interest (one antibody = one array) were incubated with the printed arrays at their pre-chosen dilution. (see SI Materials). Arrays were washed once with Zeptosens CAB1 assay buffer and incubated with primary antibodies (diluted in CAB1) over night at RT in the dark (15 h). Arrays were washed again once and incubated with Alexa647-labeled anti-species secondary antibodies (see SI Materials) for 45 min at RT in the dark. Arrays were flushed with assay buffer and imaged in the red laser of the ZeptoREADER imager system (Zeptosens). Typically, 6 images were taken at exposure times between 0.25 and 16 seconds. Non-specific assay signal contributions were evaluated from blank assays (arrays only incubated with secondary antibody). A separate protein stain assay was performed on one chip out of the print series to analyze printed protein per spot, used for normalizing the assay signals. Mean background-corrected spot signals from one image/exposure time per assay - at highest signals below saturation - were quantified, using the ZeptoVIEW 3.1 array analysis software. Signals of the two replicate samples were averaged and normalized to printed protein (NFI = normalized mean fluorescence intensity). Array/assay quality and robustness were good as verified by replicate measurements of the standard assays (phospho-Erk1/2 and phospho-EGFR). NFI signals of the co-printed control and treatment standard lysates were quantified in two independent experimental runs, on two different arrays on different chips, at different times. NFI signals and TR (treatment-to-control ratios) are given in the Table below. CVs of the NFI and of the calculated TR were in arrange below 10%. TR met their
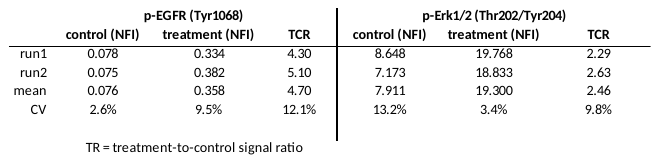
pre-defined quality criteria.

SI Materials

**Immunohistochemistry primary antibodies, their source and the working concentrations.**

| Antibody | Species | Supplier | Dilution (IHC-P) | Dilution (IHC-Fr) | Control tissue |
| --- | --- | --- | --- | --- | --- |
| monoclonal anti-human Mesothelin (420404) | rat | RD Systemouse (MAB32651) | 1:50 | 1:50 | Tonsil |
| monoclonal anti-human CA-125 (MUC16) (X325) | mouse | Abcam (ab10033) | 1:1000 | 1:1000 | Bronchus |
| monoclonal anti-human PDL1 (E1L3N) | rabbit | Cell Signaling (13684) | 1:200 | 1:200 | Tonsil |
| monoclonal anti-human CD163 (D6U1J) | rabbit | Cell Signaling (93498) | 1:500 | 1:500 | Thymus |
| polyclonal anti-human Collagen alpha-1(I) chain | rabbit | NSJ Bioreagents (R31258) | 1:1000 | 1:1000 | Skin |
| polyclonal anti-human FAP alpha | rabbit | Bio-Rad Laboratories (AHP1322) | 1:50 | 1:50 | Bronchus |
| polyclonal anti-human C1QBP | rabbit | Sigma Aldrich (A63845) | 1:50 | 1:50 | Bronchus |

**Secondary antibodies used in Immunohistochemistry and RPPA.**

| Antibody | Conjugate | IgG-type | Species | Supplier | Dilution (IHC) | Dilution (RPPA) |
| --- | --- | --- | --- | --- | --- | --- |
| anti-mouse IgG | Biotin-SP | F(ab')₂ | Goat | Jackson ImmunoResearch (115-066-062) | 1:1000 | - |
| anti-rabbit IgG | Biotin-SP | IgG (H+L) | Goat | Jackson ImmunoReseach  (111-065-144) | 1:1000 | - |
| anti-rat IgG | Biotin-SP | IgG (H+L) | Goat | Jackson ImmunoReseach  (112-065-167) | 1:1000 | - |
| anti-goat IgG | Alexa647 | F(ab')₂ | Rabbit | Invitrogen  (Z25608) | - | 1:500 |
| anti-mouse IgG | Alexa647 | IgG (H+L) | Goat | Jackson ImmunoResearch (115-605-062) | - | 1:3000 |
| anti-rabbit IgG | Alexa647 | IgG (H+L) | Goat | Invitrogen  (A21245) | - | 1:1000 |
| anti-rat IgG | Alexa647 | IgG (H+L) | Goat | Invitrogen  (A21247) | - | 1:1000 |

**Antibodies used in protein quantification by RPPA.**

| Antigen | Modification-Site | Supplier | Product No. # | Species | Dilution | Uniprot # (human) |
| --- | --- | --- | --- | --- | --- | --- |
| 4E-BP1 |  | abcam (Epitomics) | ab32024 | rabbit | 200 | Q13541 |
| 4E-BP1 - phospho | Ser65 | Cell Signaling | 9456 | rabbit | 100 | Q13541 |
| Actin beta |  | Cell Signaling | 4970 | rabbit | 5000 | P60709 |
| AFP |  | abcam (Epitomics) | ab52940 | rabbit | 200 | P02771 |
| Akt |  | Cell Signaling | 4685 | rabbit | 200 | P31749 |
| Akt - phospho | Ser473 | Cell Signaling | 4060 | rabbit | 100 | P31749 |
| Akt - phospho | Thr308 | Cell Signaling | 9275 | rabbit | 100 | P31749 |
| Aurora A (AIK) |  | Cell Signaling | 4718 | rabbit | 200 | O14965 |
| Aurora A (AIK) - phospho | Thr288 | Cell Signaling | 3079 | rabbit | 100 | O14965 |
| Aurora A/B/C - phospho | Thr288/Thr232/Thr198 | Cell Signaling | 2914 | rabbit | 100 | O14965, Q96GD4, Q9UQB9 |
| beta-Catenin |  | Millipore | 06-734 | rabbit | 500 | P35222 |
| beta-Catenin - delta |  | Cell Signaling | 59854 | rabbit | 200 | P30999 |
| beta-Catenin - phospho | Ser675 | Cell Signaling | 9567 | rabbit | 200 | P35222 |
| beta-Catenin - phospho | Thr41/Ser45 | Cell Signaling | 9565 | rabbit | 100 | P35222 |
| beta-Catenin - phospho | Ser45 | Cell Signaling | 9564 | rabbit | 100 | P35222 |
| Caspase 3 |  | Cell Signaling | 9662 | rabbit | 200 | P42574 |
| Caspase 3 - cleaved | Asp175 | Cell Signaling | 9661 | rabbit | 100 | P42574 |
| Caspase 7 - cleaved | Asp198 | abcam | ab2323 | rabbit | 100 | P55210 |
| Caspase 8 - cleaved | Asp374/Asp391 | Cell Signaling | 9496 | rabbit | 100 | Q14790 |
| CD137 |  | abcam | ab209256 | rabbit | 200 | Q07011 |
| CD163 |  | abcam | ab199427 | rabbit | 200 | Q86VB7 |
| CD44 |  | abcam (Epitomics) | ab51037 | rabbit | 200 | P16070 |
| CD44 variant (Exon v6) |  | Bender MedSystemouse | BMOUSE125 | mouse | 200 | P16070 |
| cdc2 (CDK1) |  | Cell Signaling | 9112 | rabbit | 200 | P06493 |
| cdc2 (CDK1) - phospho | Tyr15 | Cell Signaling | 4539 | rabbit | 200 | P06493 |
| CDK2 |  | Cell Signaling | 2546 | rabbit | 200 | P24941 |
| CDK2 - phospho | Thr160 | Cell Signaling | 2561 | rabbit | 100 | P24941 |
| CDK4 |  | Cell Signaling | 12790 | rabbit | 200 | P11802 |
| CDK4 - phospho | Thr172 | Invitrogen | PA-64482 | rabbit | 100 | P11802 |
| CDK6 |  | Cell Signaling | 13331 | rabbit | 200 | Q00534 |
| CDK6 - phospho | Tyr24 | biorabbityt | orabbit15014 | rabbit | 100 | Q00534 |
| c-Jun |  | Cell Signaling | 9165 | rabbit | 200 | P05412 |
| c-Jun - phospho | Ser73 | Cell Signaling | 9164 | rabbit | 100 | P05412 |
| c-Jun - phospho | Ser63 | Cell Signaling | 2361 | rabbit | 100 | P05412 |
| c-Met (HGF/SF receptor) - phospho | Tyr1349 | abcam (Epitomics) | ab68141 | rabbit | 100 | P08581 |
| c-Raf |  | Cell Signaling | 9422 | rabbit | 200 | P04049 |
| c-Raf - phospho | Ser259 | Cell Signaling | 9421 | rabbit | 100 | P04049 |
| Cyclin B1 |  | abcam | ab32053 | rabbit | 200 | P14635 |
| Cyclin B1 - phospho | Ser133 | Cell Signaling | 4133 | rabbit | 100 | P14635 |
| Cyclin E2 |  | abcam (Epitomics) | ab32103 | rabbit | 200 | O96020 |
| E-Cadherin |  | R&D | AF748 | goat | 200 | P12830 |
| EGFR (Erabbit-1, HER1) |  | Cell Signaling | 4405 | rabbit | 200 | P00533 |
| EGFR (Erabbit-1, HER1) - phospho | Tyr1068 | Cell Signaling | 2234 | rabbit | 100 | P00533 |
| EGFR (Erabbit-1, HER1) - phospho | Tyr845 | Cell Signaling | 2231 | rabbit | 100 | P00533 |
| EGFR (Erabbit-1, HER1) - phospho | Tyr992 | Cell Signaling | 2235 | rabbit | 100 | P00533 |
| EGFR (Erabbit-1, HER1) - phospho | Tyr1068 | Cell Signaling | 2234 | rabbit | 100 | P00533 |
| EGFR (Erabbit-1, HER1) - phospho | Tyr1068 | Cell Signaling | 2234 | rabbit | 100 | P00533 |
| eIF2 alpha |  | Cell Signaling | 9722 | rabbit | 200 | P05198 |
| eIF2 alpha - phospho | Ser51 | Cell Signaling | 3398 | rabbit | 100 | P05198 |
| eIF4E |  | Cell Signaling | 2067 | rabbit | 200 | P06730 |
| eIF4E - phospho | Ser209 | Cell Signaling | 9741 | rabbit | 100 | P06730 |
| EpCAM (CD326) |  | Cell Signaling | 3599 | rabbit | 200 | P16422 |
| Erk1/2 (MAPK p44/42) |  | Cell Signaling | 4695 | rabbit | 200 | P27361, P28482 |
| Erk1/2 (MAPK p44/42) - phospho | Thr202/Tyr204 | Cell Signaling | 4370 | rabbit | 100 | P27361, P28482 |
| Erk1/2 (MAPK p44/42) - phospho | Thr202/Tyr204 | Cell Signaling | 4370 | rabbit | 100 | P27361, P28482 |
| Erk1/2 (MAPK p44/42) - phospho | Thr202/Tyr204 | Cell Signaling | 4370 | rabbit | 100 | P27361, P28482 |
| FAK1 - phospho | Tyr397 | Cell Signaling | 8556 | rabbit | 100 | Q05397 |
| FGF receptor 1 |  | Cell Signaling | 9740 | rabbit | 200 | P11362 |
| GSK3 alpha/beta - phospho | Ser21/Ser9 | Cell Signaling | 9331 | rabbit | 100 | P49840 , P49841 |
| GSK3 alpha/beta - phospho | Tyr279/Tyr216 | abcam | ab68476 | rabbit | 100 | P49840 , P49841 |
| GSK3 beta |  | Cell Signaling | 9315 | rabbit | 200 | P49841 |
| GSK3 beta - phospho | Ser9 | Cell Signaling | 9336 | rabbit | 100 | P49841 |
| Her2 |  | Dako | A0485 | rabbit | 200 | P04626 |
| Her2 - phospho | Tyr1248 | Cell Signaling | 2247 | rabbit | 100 | P04626 |
| Histone H3 |  | Cell Signaling | 9715 | rabbit | 10000 | P68431 |
| Histone H3 - acetyl | Lys9/Lys14 | Calbiochem | 382158 | rabbit | 2000 | P68431 |
| Histone H3 - phospho | Ser10 | Cell Signaling | 9701 | rabbit | 100 | P68431 |
| Histone H3 - phospho | Thr11 | Cell Signaling | 9764 | rabbit | 100 | P68431 |
| Histone H3 - trimethyl | Lys27 | Cell Signaling | 9733 | rabbit | 100 | P68431 |
| IkappaB alpha |  | Cell Signaling | 9242 | rabbit | 200 | P25963 |
| IkappaB alpha - phospho | Ser32 | Cell Signaling | 9241 | rabbit | 100 | P25963 |
| Ki-67 |  | Dako | M7240 | mouse | 200 | P46013 |
| MEK1 |  | Cell Signaling | 9124 | rabbit | 200 | Q02750 |
| MEK1/2 - phospho | Ser217/Ser221 | Cell Signaling | 9154 | rabbit | 100 | Q02750, P36507 |
| MEK2 |  | Cell Signaling | 9125 | rabbit | 200 | P36507 |
| Mesothelin |  | R&D Systemouse | MAB32651 | rat | 200 | Q13421 |
| MKK4 (SEK1) - phospho | Ser257/Thr261 | Cell Signaling | 9156 | rabbit | 100 | P45985 |
| mTOR (FRAP) |  | Cell Signaling | 2983 | rabbit | 200 | P42345 |
| mTOR (FRAP)- phospho | Ser2448 | Cell Signaling | 2971 | rabbit | 100 | P42345 |
| Nanog |  | Cell Signaling | 4903 | rabbit | 200 | Q9H9S0 |
| N-Cadherin |  | BD Transduction | 610920 | mouse | 200 | P19022 |
| NF-κB p65 |  | abcam (Epitomics) | ab76311 | rabbit | 200 | Q04206 |
| NF-κB p65 - phospho | Ser536 | Cell Signaling | 3033 | rabbit | 100 | Q04206 |
| Oct-4 |  | Cell Signaling | 4286 | mouse | 200 | Q01860 |
| p27 (Kip1, CDKN1B) - phospho | Ser10 | abcam (Epitomics) | ab62364 | rabbit | 100 | P46527 |
| p38 MAPK |  | Cell Signaling | 9212 | rabbit | 200 | Q16539 |
| p38 MAPK - phospho | Thr180/Tyr182 | Cell Signaling | 9211 | rabbit | 100 | Q16539 |
| p53 |  | Cell Signaling | 2527 | rabbit | 200 | P04637 |
| p53 - acetyl | Lys382 | Cell Signaling | 2525 | rabbit | 100 | P04637 |
| p70 S6 Kinase |  | Cell Signaling | 2708 | rabbit | 200 | P23443 |
| p70 S6 kinase - phospho | Thr421/Ser424 | Cell Signaling | 9204 | rabbit | 100 | P23443 |
| p70 S6 Kinase - phospho | Thr389 | Cell Signaling | 9205 | rabbit | 100 | P23443 |
| PARP |  | Cell Signaling | 9532 | rabbit | 200 | P09874 |
| PARP - cleaved | Asp214 | Cell Signaling | 9541 | rabbit | 100 | P09874 |
| PCNA |  | Kremmer |  | rat | 200 | P12004 |
| PD1 |  | Cell Signaling | 86163 | rabbit | 200 | Q15116 |
| PDGF receptor beta |  | Cell Signaling | 3169 | rabbit | 200 | P09619 |
| PD-L1 |  | Cell Signaling | 13684 | rabbit | 200 | Q9NZQ7 |
| PI3-kinase p110 beta |  | Millipore | 04-400 | rabbit | 200 | P42338 |
| PI3-kinase p85 alpha |  | abcam (Epitomics) | ab40755 | rabbit | 200 | P27986 |
| PP1 alpha - phospho | Thr320 | abcam (Epitomics) | ab62334 | rabbit | 100 | P62136 |
| PTEN |  | Cell Signaling | 9552 | rabbit | 200 | P60484 |
| PTEN - phospho | Ser380 | Cell Signaling | 9551 | rabbit | 100 | P60484 |
| Rad51 |  | abcam (Epitomics) | ab109107 | rabbit | 200 | Q06609 |
| Rabbit |  | Cell Signaling | 9313 | rabbit | 200 | P06400 |
| Rabbit - phospho | Ser807/Ser811 | Cell Signaling | 8516 | rabbit | 100 | P06400 |
| RSK 1 (p90RSK) - phospho | Ser380 | Cell Signaling | 9341 | rabbit | 100 | Q15418 |
| RSK 1 (p90RSK) - phospho | Thr573 | abcam (Epitomics) | ab62324 | rabbit | 100 | Q15418 |
| RSK 1/2/3 |  | Cell Signaling | 9347 | rabbit | 200 | Q15418, Q51812, Q15349 |
| RSK 3 - phospho | Thr356/Ser360 | Cell Signaling | 9348 | rabbit | 100 | Q15349 |
| S6 ribosomal protein |  | Cell Signaling | 2217 | rabbit | 200 | P62753 |
| S6 ribosomal protein - phospho | Ser240/Ser244 | Cell Signaling | 2215 | rabbit | 100 | P62753 |
| S6 ribosomal protein - phospho | Ser235/Ser236 | Cell Signaling | 2211 | rabbit | 100 | P62753 |
| Slug |  | Cell Signaling | 9585 | rabbit | 200 | O43623 |
| Smad1 |  | Cell Signaling | 6944 | rabbit | 200 | Q15797 |
| Smad2 - phospho | Ser465/Ser467 | Cell Signaling | 3108 | rabbit | 100 | Q15796 |
| Smad2 - phospho | Ser245/Ser250/Ser255 | Cell Signaling | 3104 | rabbit | 100 | Q15796 |
| Smad2/3 |  | Cell Signaling | 3102 | rabbit | 200 | Q15796, P84022 |
| Snail |  | Cell Signaling | 3879 | rabbit | 200 | O95863 |
| Src |  | Cell Signaling | 2108 | rabbit | 200 | P12931 |
| Src - phospho | Tyr527 | Cell Signaling | 2105 | rabbit | 100 | P12931 |
| STAT 1 |  | Cell Signaling | 9175 | rabbit | 200 | P42224 |
| STAT 1 - phospho | Tyr701 | Cell Signaling | 9167 | rabbit | 100 | P42224 |
| STAT 3 |  | Cell Signaling | 4904 | rabbit | 200 | P40763 |
| STAT 3 - phospho | Ser727 | Cell Signaling | 9134 | rabbit | 100 | P40763 |
| STAT 3 - phospho | Tyr705 | Cell Signaling | 9145 | rabbit | 100 | P40763 |
| Tubulin acetylated |  | Sigma | T6793 | mouse | 200 | P68366 |
| WT1 (Wilms Tumor 1) |  | Cell Signaling | 83535 | rabbit | 200 | P19544 |

**FACS antibodies, their source and the working concentrations.**

| Antibody | Species | Supplier | Staining | Panel | Dilution |
| --- | --- | --- | --- | --- | --- |
| anti-human CD3, FITC | mouse | BD Biosciences (345763) | extracellular | 1, 2, 5 | 1:40 |
| anti-human CD8, PerCP | mouse | Thermo Fisher Scientific (MHC0831) | extracellular | 1, 2 | 1:40 |
| anti-human CD8, FITC | mouse | BioLegend  (301005) | extracellular | 4, 7 | 1:20 |
| anti-human CD4, PE-Cy7 | mouse | BioLegend  (344611) | extracellular | 1, 3 | 1:20 |
| anti-human CD137, PE | mouse | Miltenyi Biotec  (130-098-878) | extracellular | 2 | 1:11 |
| anti-human CD137, BV421 | mouse | BioLegend  (309820) | extracellular | 1 | 1:20 |
| anti-human CD25, PerCP | mouse | BioLegend  (356131) | extracellular | 3 | 1:20 |
| anti-human Foxp3, BV421 | mouse | BioLegend  (320123) | intracellular | 3 | 1:20 |
| anti-human CD39, PE-Dazzle 594 | mouse | BioLegend  (328223) | extracellular | 4 | 1:20 |
| anti-human PD1, BV421 | mouse | BioLegend  (329919) | extracellular | 4, 7 | 1:20 |
| anti-human CTLA4, PerCP-Cy5.5 | mouse | BioLegend  (369607) | extracellular | 4 | 1:20 |

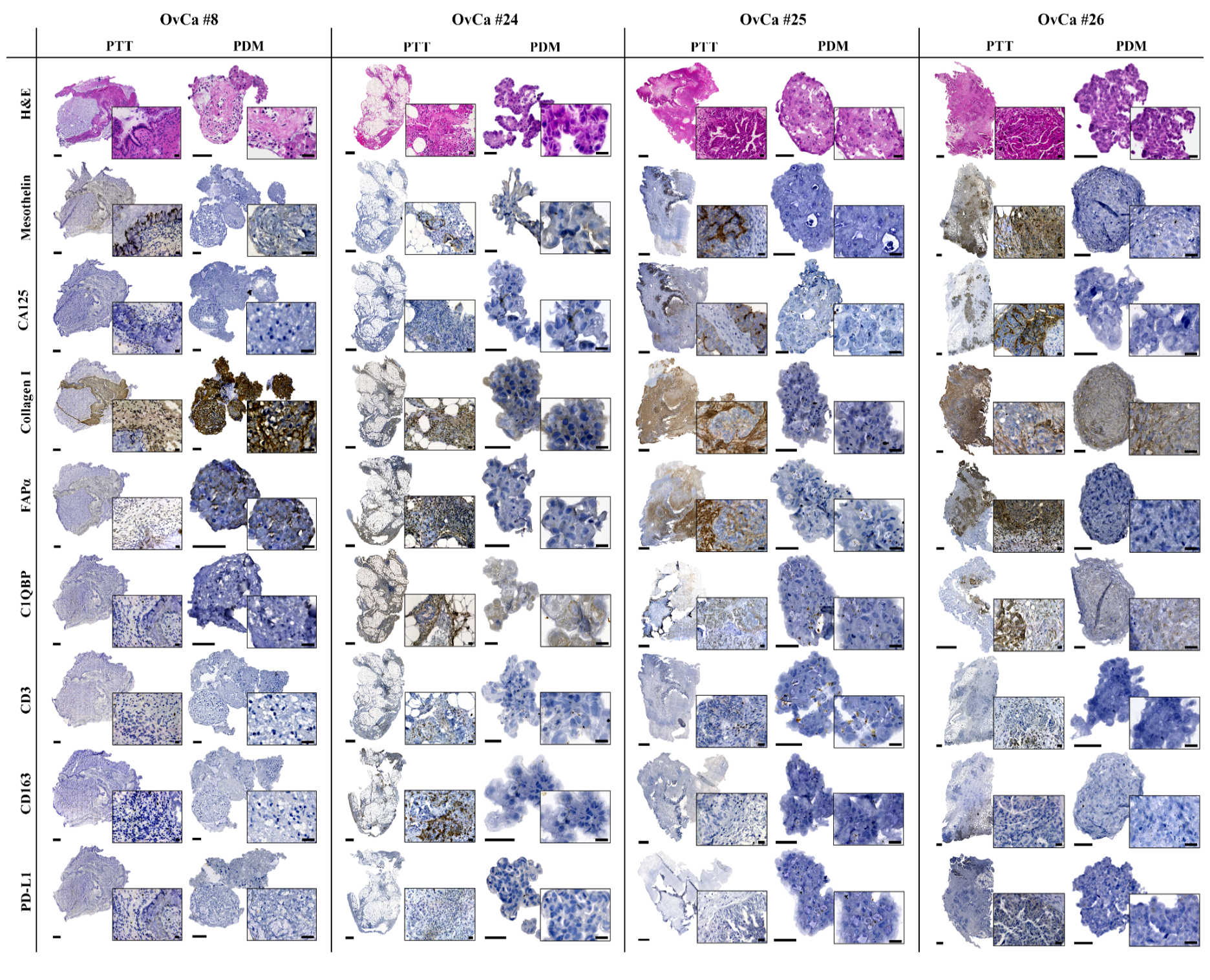

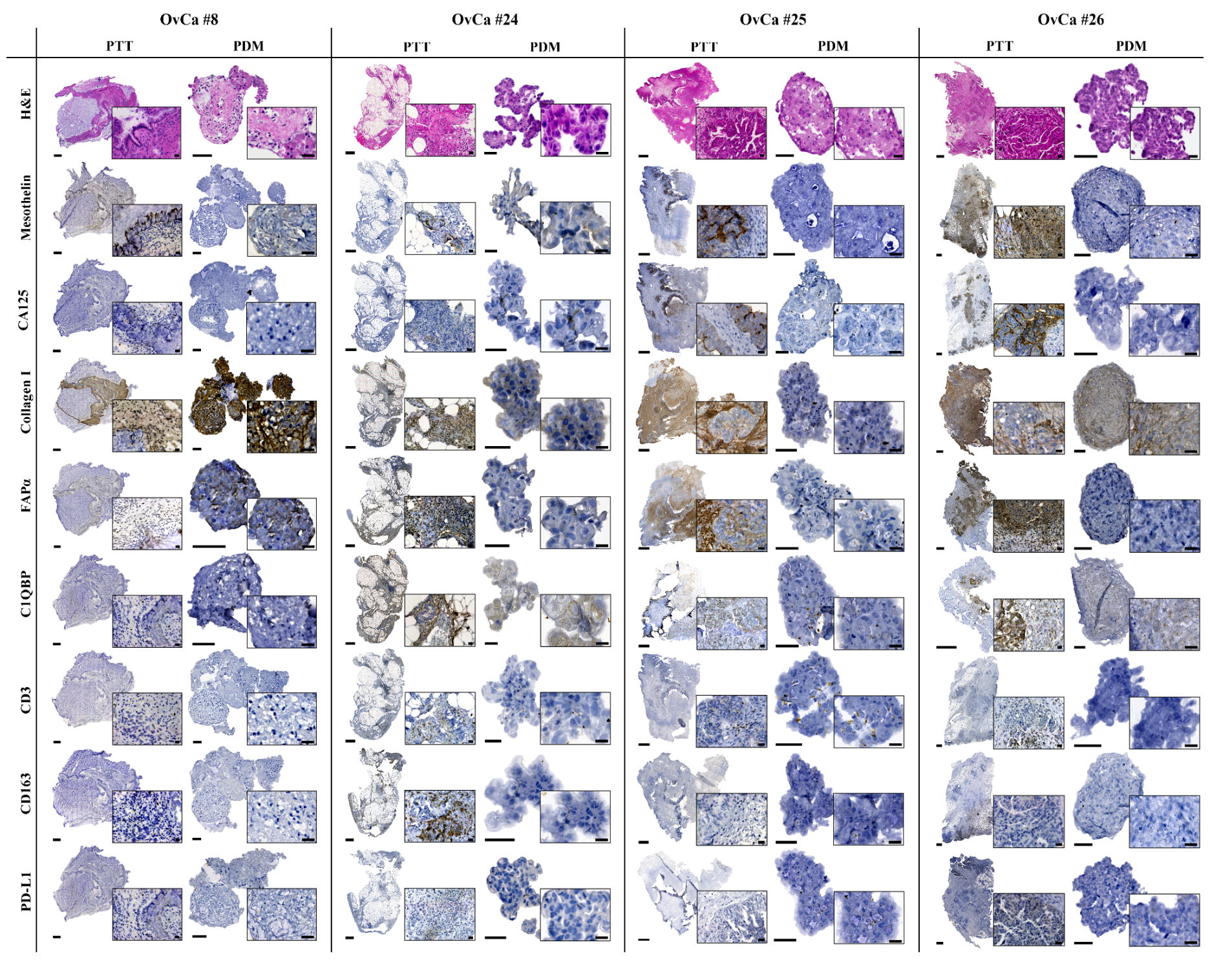

**Fig. S1.** Histology and immunohistochemistry of OvCa microtumors and corresponding primary tumor tissue. Hematoxylin and Eosin (H&E) staining of OvCa patient-derived microtumors (PDM) and corresponding primary tumor tissue (PTT) revealed several features of malignant cells. Qualitative characterization of OvCa microtumors and PTT by DAB immunohisto-chemical staining. Expression of tumor markers (CA125, mesothelin), tumor-associated macrophages CD163), immune/tumor marker (PD-L1), cancer-associated fibroblasts (FAPα) and extracellular matrix (Hyaloronan C1QBP, Collagen I) within PDMs and PTT are shown. Scale bars indicate 500 µm for PTT; 50 µm for PDM; 20 µm for magnifications (PTT and PDM).

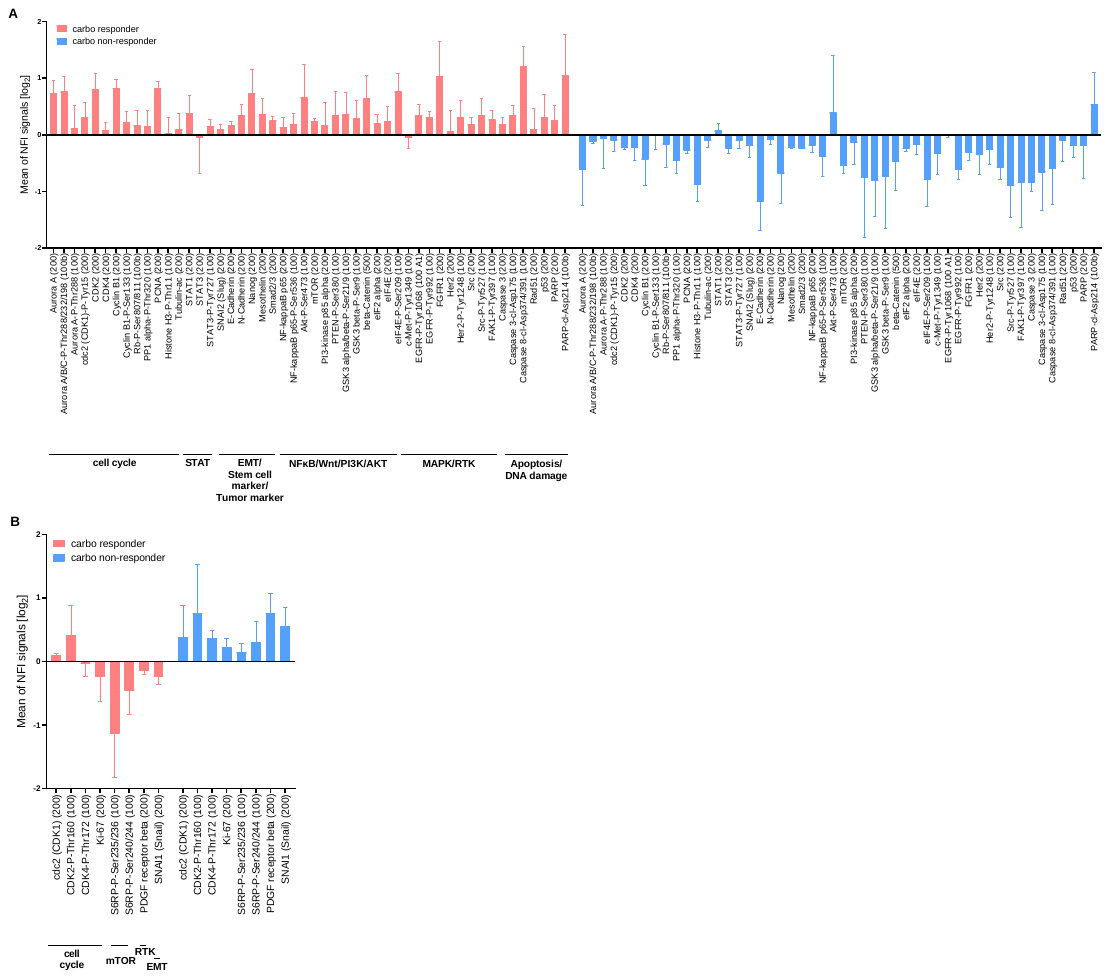

**Fig. S2**. **Up- and downregulated proteins in carboplatin responder and non-responder OvCa PDM.** Mean protein abundances (NFI) within carboplatin responder and non-responder of untreated OvCa PDM. Proteins with >20% differential mean NFI signals between responder and non-responder were selected and plotted according to pathway affiliation. Data are shown as mean ± SEM (**A**) Proteins upregulated in carboplatin responder PDM. (**B**) Proteins downregulated in carboplatin responder PDM.

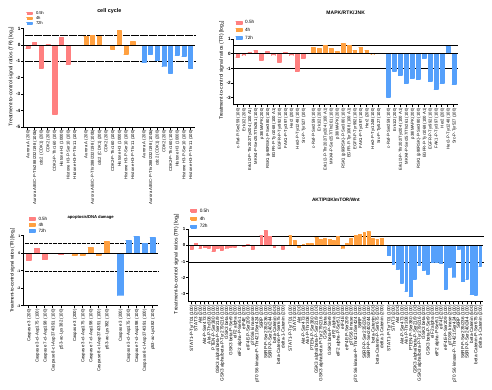

**Fig. S3.** **Time-dependent alterations of signaling pathways in carboplatin-sensitive OvCa PDM.** On- and off-target effects of carboplatin treatment (75 µM) were assessed by RPPA analysis of treatment sensitive OvCa PDM. PDM from OvCa #24 were treated with carbo for different time spans to examine proteomic changes in the course of treatment. Protein abundances are displayed as treatment-to-control signal ratios (TR) calculated from NFI signals of treated PDM and DMSO vehicle control for each time point, log_2_-transformed and sorted according to pathway affiliation. A threshold of >50% differential protein expression between treated samples and vehicle control was applied. Protein abundances are shown from 0.5 hours, 4 hours and 72 hours treated PDM. Carbo, carboplatin.

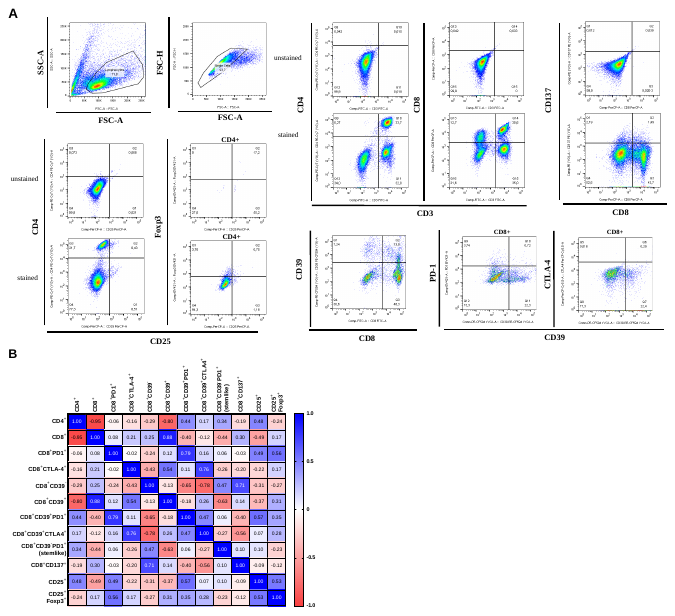

**Fig. S4. Gating schemes of expanded TILs, the correlation of TIL populations and their comparison between different OvCa models.** (**A**) Exemplary FACS plots of multicolor stained OvCa TILs (OvCa #4) measured with FACS Melody (BD Biosciences) compared to unstained controls. (**B**) Nonparametric Spearman correlation of characterized TIL populations within n = 13 OvCa TIL samples. Shown are Spearman R-values. CTL, Cytotoxic lymphocytes

Table S1. Correlation of PDM isolation-success and clinical patient data (Spearman correlation).

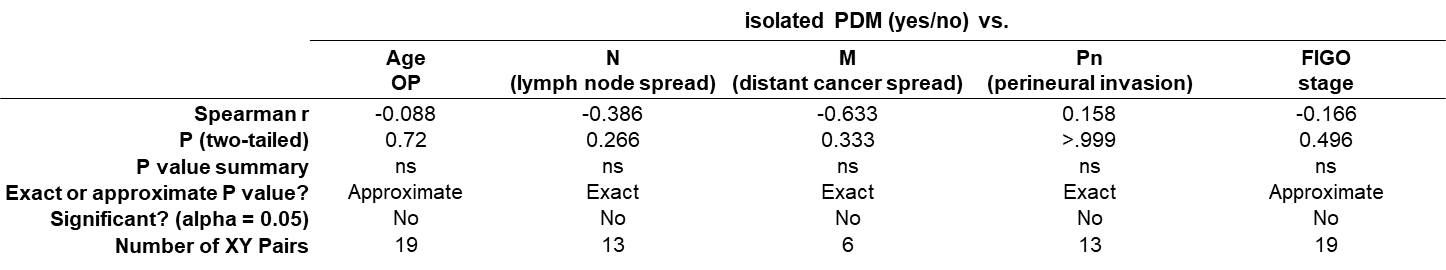

Age was ranked into < 66 years = “0” and ≥ 66 years = “1”; N-M-Pn and PDM-isolation were classified as “yes” or “no”.

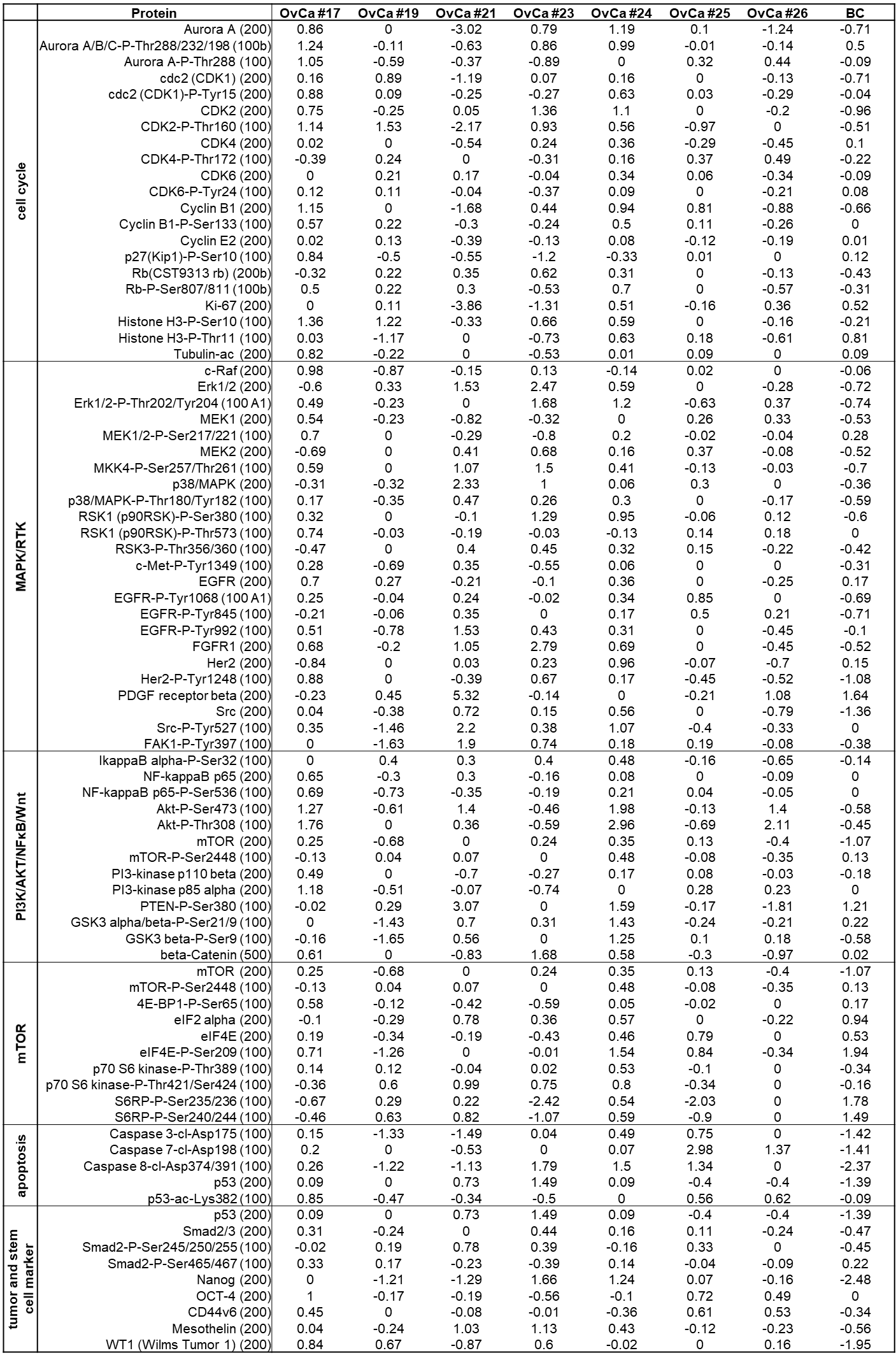
Table S2. Log_2_-transformed, median-centered NFI signals of signaling pathway proteins from OvCa and BC PDM from RPPA analysis.

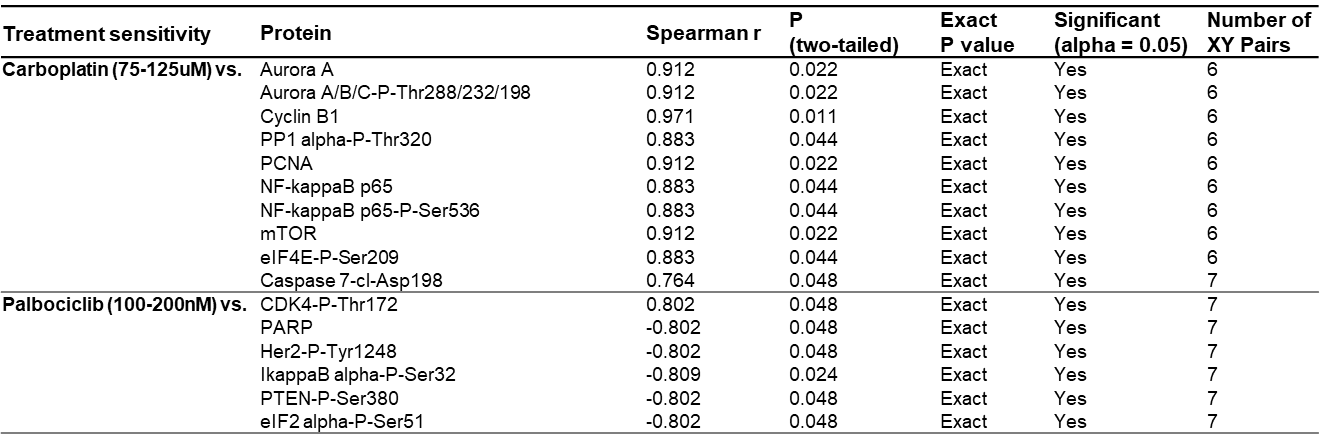
Table S3. Spearman correlation of carboplatin-treatment sensitivity and protein abundances of OvCa PDM models.

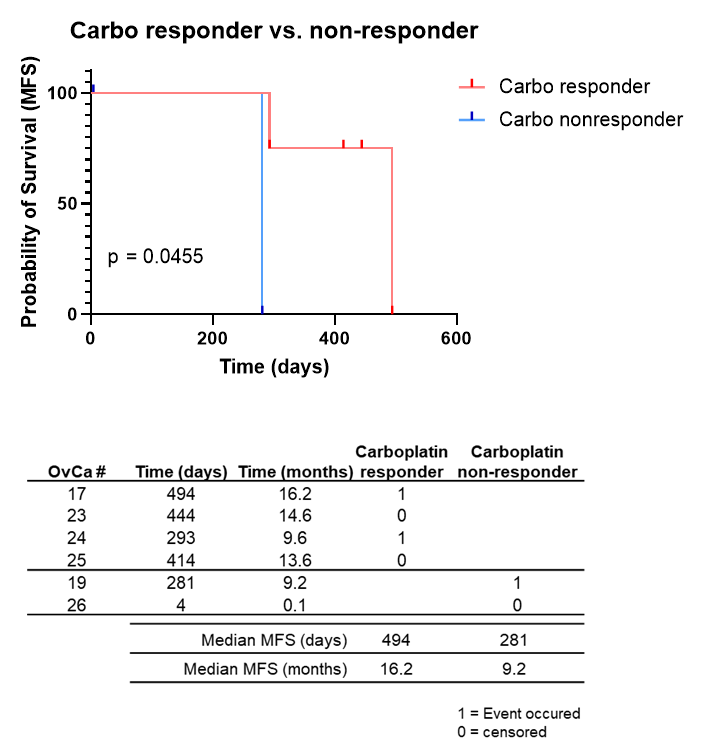

Table S4. Significant difference in metastasis-free-survival between OvCa PDM carboplatin responder and non-responder.

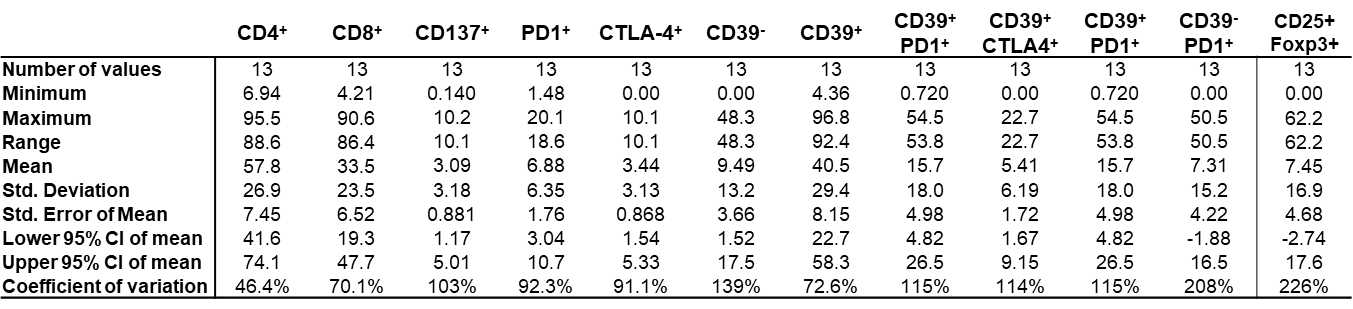
Table S5. Descriptive statistics of analyzed OvCa TIL populations.

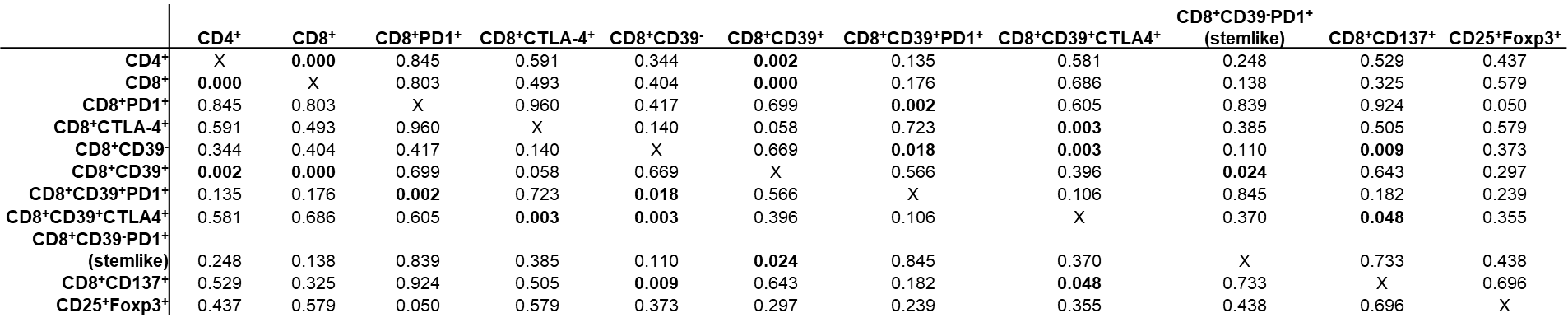
Table S6. Correlation of TIL phenotypes (Spearman correlation; P-Values).
